# 3D-Printing of Electroconductive MXene-based Micro-meshes in a Biomimetic Hyaluronic Acid-based Scaffold Directs and Enhances Electrical Stimulation for Neural Repair Applications

**DOI:** 10.1101/2024.04.05.587425

**Authors:** Ian Woods, Dahnan Spurling, Sandra Sunil, Jack Maughan, Javier Gutierrez-Gonzalez, Tara K. McGuire, Liam Leahy, Adrian Dervan, Valeria Nicolosi, Fergal J O’Brien

## Abstract

No effective treatments are currently available for central nervous system neurotrauma although recent advances in electrical stimulation suggest some promise in neural tissue repair. We hypothesized that structured integration of an electroconductive biomaterial into a tissue engineering scaffold could enhance electroactive signalling for neural regeneration.

Electroconductive 2D Ti_3_C_2_T*_x_* MXene nanosheets were synthesized from MAX-phase powder, demonstrating excellent biocompatibility with neurons, astrocytes and microglia. To achieve spatially-controlled distribution of these MXenes, melt-electrowriting was used to 3D-print highly-organized PCL micro-meshes with varying fibre spacings (low-, medium-and high-density), which were functionalized with MXenes to provide highly-tunable electroconductive properties (0.081±0.053-18.87±2.94 S/m). Embedding these electroconductive micro-meshes within a neurotrophic, immunomodulatory hyaluronic acid-based extracellular matrix (ECM) produced a soft, growth-supportive MXene-ECM composite scaffold. Electrical stimulation of neurons seeded on these scaffolds promoted neurite outgrowth, influenced by fibre spacing in the micro-mesh. In a multicellular model of cell behaviour, neurospheres stimulated for 7 days on high-density MXene-ECM scaffolds exhibited significantly increased axonal extension and neuronal differentiation, compared to low-density scaffolds and MXene-free controls. The results demonstrate that spatial-organization of electroconductive materials in a neurotrophic scaffold can enhance repair-critical responses to electrical stimulation and that these biomimetic MXene-ECM scaffolds offer a promising new approach to neurotrauma repair.

## 1. Introduction

Neurotrauma is associated with impairments of cognitive, motor and sensory function and can result in paralysis, disability and cognitive difficulties^[1]^. The complex pathophysiology of neurotrauma, including inflammation, scarring and poor neuronal regrowth, makes the development of effective treatments extremely challenging^[2,3]^ and suggests that a multi-pronged approach may be required to repair central nervous system (CNS) injuries^[4]^. Biomaterial-based strategies offer platforms which can address aspects of neural injuries through providing an axonal growth-supporting environment to act as a bridge across the injury site for neuronal reconnection while also enabling the local delivery of therapeutic strategies^[3,5]^. One such strategy, therapeutic electrical stimulation (ES), holds particular promise due to its multi-faceted therapeutic potential. Several *in vivo* studies have demonstrated the potential of exogenous ES, applied using transcutaneous or epidurally-placed electrodes, to activate isolated neuronal circuitry in the injured spinal cord as well as to encourage regrowth of injured axons when applied to neuronal cell bodies in the brain^[6,7]^. While encouraging, these approaches have the potential to target only certain subsets of affected injured neurons. Accumulating *in vitro* evidence suggests that ES applied directly to axons can promote their growth^[8]^ while also enhancing neuronal plasticity^[9]^ and supporting neuronal function^[10]^ and has also been shown to activate key cell signalling pathways in neurons, such as the mammalian target of rapamycin (mTor) ^[11,12]^ - a key regulator of axonal growth^[13]^.

The electroconductivity of a biomaterial plays a key role in determining how electric fields interact with neural tissues in tissue engineering scaffolds and 3D environments^[14]^. For example, the delivery of ES to neurons via conductive biomaterial interfaces has been shown to enhance the differentiation of neuronal progenitor cells, promote axonal growth and even enhance synaptic plasticity^[9,15–18]^. Previous work by Leahy et al. (2024) has shown that a conductive polypyrrole-coated PCL axonal tract geometry, based on the anatomical organization of spinal cord tracts, can enhance *in vitro* neurite outgrowth from neurons in response to ES^[19]^. *In vivo*, the incorporation of an electroactive sheath within tissue engineering conduits for peripheral nerve repair has been used to promote functional recovery in animal models of sciatic nerve injury^[20]^.These studies raise the question of how the design of electroactive tissue engineering scaffolds can be optimized to enhance the therapeutic properties of electrical stimulation and how electroconductive biomaterials should be integrated into tissue engineering scaffolds to facilitate such strategies. Furthermore, the specific spatial distribution of electroconductive materials in tissue engineering scaffold design remains poorly investigated.

The most commonly used biomaterials for the therapeutic delivery of ES are metallic electrodes – often used for deep brain stimulation treatments (e.g. for Parkinson’s Disease)^[21]^. The electrochemical properties of metals are excellently suited for sensor and stimulation applications but these stiff materials exhibit significant limitations including compliance mismatch with neural tissues^[22]^, poor neurocompatibility^[23]^ slow degradation rates, and the release of potentially toxic ions thus making them unsuitable for neural tissue engineering scaffold design^[24–27]^. In order to overcome these issues, a range of softer and more biomimetic conductive biomaterials such as conductive polymers and nanocomposite biomaterials have been developed^[28,29]^. However, conductive polymer-based biomaterials generally exhibit three key limitations – poor processability or incompatibility with additive manufacturing technologies^[30,31]^, limited bioactivity^[32]^ and rapid degradation of electroconductive properties in biological environments^[33]^. The incorporation of conductive nanomaterials within natural (e.g. collagen, alginate)^[29,34]^ and synthetic (e.g. PCL)^[35]^ polymers can form composites that are biocompatible and can be used to produce highly processable materials, although high volume ratios are often required to achieve percolation of the nanomaterial phase within natural polymers in order to form an electrically conductive bulk material, leading to high stiffness and low bioactivity^[36]^. If these limitations can be overcome, the processability of these nanocomposite biomaterials opens new opportunities for the design of sophisticated neurocompatible tissue engineering scaffolds. One promising class of bio-nanomaterials are 2D materials such as MXenes, specifically Ti_3_C_2_T*_x_* nanosheets derived from the layered ceramic Ti_3_AlC_2_ MAX phase which exhibit excellent stability, high conductivity and broad biocompatibility^[17,37]^.

We hypothesized that the structured integration of a MXene nanosheet network within an extracellular matrix-based (ECM) tissue engineering scaffold, with proven potential for neural applications, would produce an electroconductive scaffold capable of enhancing delivery of electrical stimulation to ingrowing neurons. First, the electroconductive properties of Ti_3_C_2_T_x_ MXene substrates was measured and their biocompatibility tested with key neural cell types (neurons, astrocytes and microglia). Using melt electrowriting, we initially manufactured polycaprolactone micro-meshes of varying microfibre densities and then functionalized them with the MXene nanosheets. These electroconductive MXene/PCL micro-meshes were then embedded within a biomimetic neurotrophic macroporous extracellular matrix-based (ECM) scaffold consisting of hyaluronic acid (HA), collagen type-IV and fibronectin which we previously developed as a neurotrophic, immunomodulatory platform for spinal cord injury (SCI) repair - in which collagen-IV and fibronectin synergistically enhanced axonal growth and pro-reparative astrocyte signalling in a stiffness-dependent manner^[38,39]^. These biphasic composite MXene-ECM scaffolds provided a soft neurotrophic environment to enhance axonal growth and, when subjected to electrical stimulation, the MXene-ECM scaffolds enhanced neurite outgrowth and this behaviour was significantly affected by micro-mesh design. To examine these effects in a multicellular model of cell behaviour, murine olfactory bulb-derived neurospheres were subjected to electrical stimulation for 7 days resulting in enhanced axonal growth and neuronal differentiation on high density MXene micro-meshes compared to those cultured on MXene-free scaffold controls.

## 2. Results

### 2.1 MXene synthesis, characterization and MXene-functionalisation of PCL substrates

Ti_3_C_2_T*_x_* MXenes are highly electrically conductive 2D nanosheets which exhibit excellent biocompatibility^[40]^. To produce a MXene ink for fabrication of electroconductive films, multilayer Ti_3_C_2_T*_x_* MXenes were etched from Al-rich Ti_3_AlC_2_ MAX phase powder to take advantage of the excellent degradation resistance afforded by isolating MXenes from a MAX phase precursor with higher Al-content^[41]^. These multilayer stacks were then dispersed into delaminated sheets using vortex mixing and then centrifuged to form a concentrated MXene ink in an aqueous solution. SEM analysis (Figure 1a) of the resulting MXene flakes reveal the successful synthesis of 2D nanosheets with a micrometre scale diameter. EDX analysis (Figure 1b) of the elemental composition of the flakes indicates the distribution of characteristic functional groups of Ti_3_C_2_T*_x_* MXenes, such as C-, Ti-and O-groups, and the average flake diameter (Figure 1c) was approximately 1.87 ± 1.18 μm. SEM analysis of vacuum-annealed MXene films (Figure 1d) demonstrated the nanoscale thinness of the sheets, their high aspect ratio and the multi-layered morphology of MXene films. Analysis of the conductivity (Figure 1e) of annealed MXene films indicated that the layered 2D nanosheets exhibited a high conductivity of approximately 10,800 ± 200 S/cm. However, while Ti_3_C_2_T*_x_* MXene films exhibit excellent electroconductivity, they also possess poor mechanical integrity in the absence of a supporting material. To produce robust MXene sheet networks, MXene films were templated across PCL substrates (Figure 1f) to produce MXene/PCL bilayered films. To improve the adhesion of the MXene nanosheets to the PCL surface, the PCL films were first treated with NAOH^[42,43]^ to improve the hydrophilicity of the film surface (Figure 1g). AFM analysis of the NAOH-treated PCL films indicate a pitted polymer surface (Figure 1h) while the MXene coating formed a scale-like coating along the fibre surface (Figure 1i). In high-resolution phase-based imaging of the surface (Figure 1j) the layered flake morphology of the intercalated MXene nanosheet coatings can be clearly observed. These results demonstrate that the surface charge interactions between NAOH-treated PCL surfaces and hydrophilic MXene nanosheets result in the formation of a cohesive MXene film on the PCL surface, producing a MXene-PCL bi-layered film. This approach combines the excellent processability of PCL with the high electroconductivity of MXene nanosheets, producing an electroconductive biomaterial compatible with a wide range of biofabrication approaches - including 3D-printing and melt-electrowriting.

**Figure 1.**
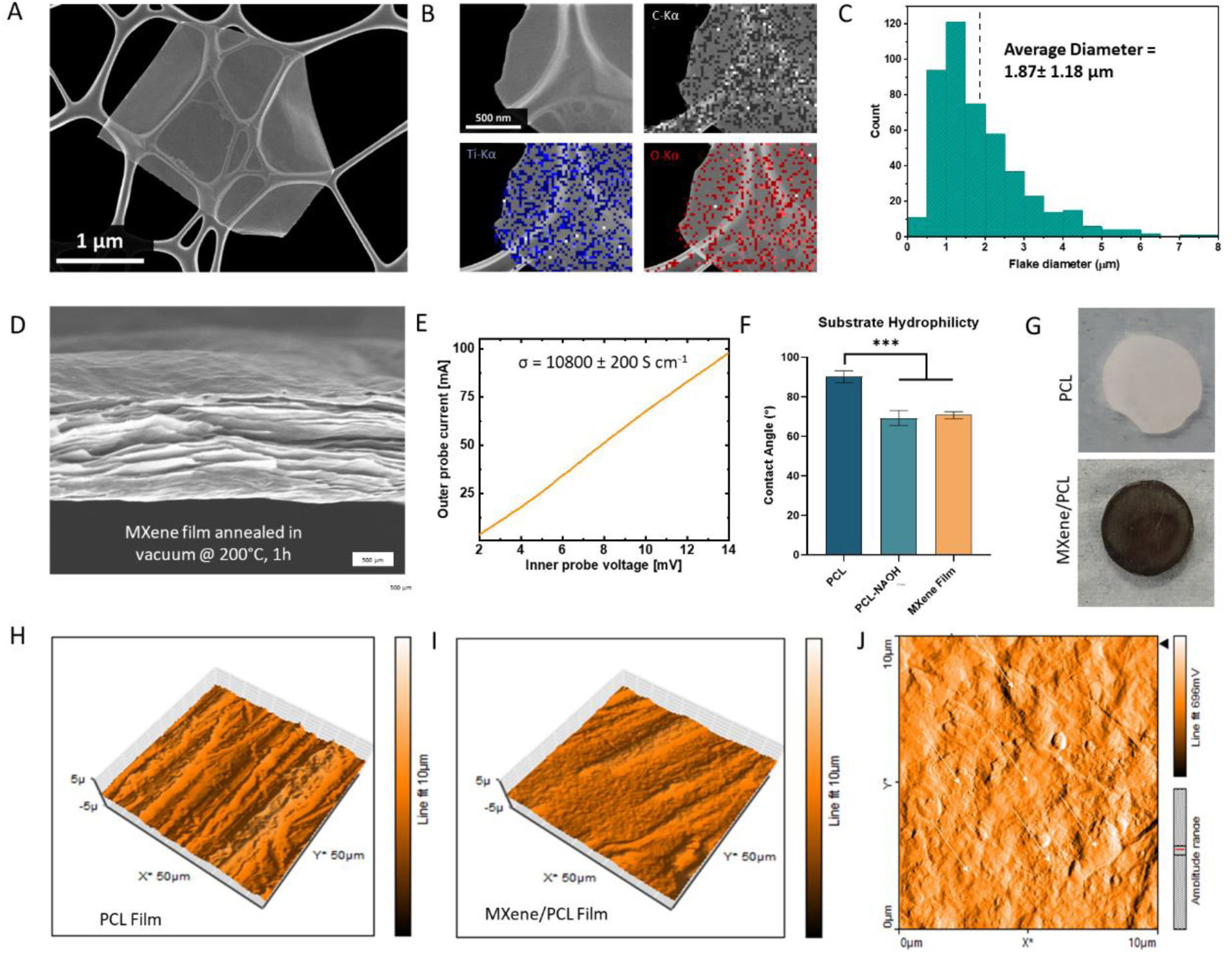
MXene and MXene-PCL Characterization. (A) SEM micrograph of MXene flake morphology and (B) EDX analysis of elemental composition. (C) Nanoparticle size distribution measurements, indicating an average particle diameter of 1.87 ± 1.18 μm. (D) SEM of vacuum-annealed MXene film, demonstrating layered flake morphology. (E) Conductivity of annealed MXene films. (F) NAOH-treated PCL surfaces exhibit similar hydrophilicity to MXene coated films. (G) Coating of PCL films produces opaque black film. Atomic Force Microscopy of tip-displacement analysis (H) PCL and (I) MXene-PCL films. (J) High-resolution deflection measurement of MXene-PCL surfaces, white arrows indicate raised single MXene sheets. *** indicates p <0.001, p values calculated using ANOVA with Tukey post-hoc testing, error bars indicate S.E.M.

### 2.2 Neurocompatibility of MXene-functionalized PCL surfaces

To assess the biocompatibility of the MXene/PCL bi-layered films with key neural cell types, PCL films with, or without, MXene coatings were seeded with SH-SY5Y human neuronal cells, murine IMG microglial cells and human-derived astrocytes (Figure 2). All cell types exhibited widespread coverage on both the PCL and MXene/PCL films with confluent monolayers formed across both samples (Figure 2a). Neurons extended neurites across both surfaces, indicative of cytocompatibility. Astrocytes and microglia exhibited excellent coverage on the MXene/PCL substrates and astrocytes exhibited non-hypertrophic morphologies, extending long processes. Neurons displayed significantly increased proliferation (Figure 2b) on the MXene/PCL substrates, which was matched by a 4.51 ± 1.02-fold change (p<0.001) increase in metabolic activity (Figure 2c). Similarly, astrocytes adhered excellently to the MXene/PCL/ substrates and showed increased metabolic activity at day 4 (Figure 2d, e), but lower expression of glial acidic fibrillary protein (GFAP) - a marker of reactivity in astrocytes (Figure 2f, p<0.05). Finally, although no significant difference between PCL and MXene/PCL substrates on microglial proliferation or metabolic activity was observed, microglia did proliferate across both substrates equally (Figure 2g, h), indicative of low cytotoxicity. Together, these results suggest that MXene-coated PCL substrates provide a neurocompatible biomaterial interface for key neural cell types. Furthermore, the increased neuronal and astrocyte proliferation, alongside decreased expression of GFAP, suggests that the addition of the MXene coating improves the neurocompatibility of the coated substrate beyond that of the PCL-only controls – one of the most commonly used biopolymers for tissue engineering applications^[44]^.

**Figure 2.**
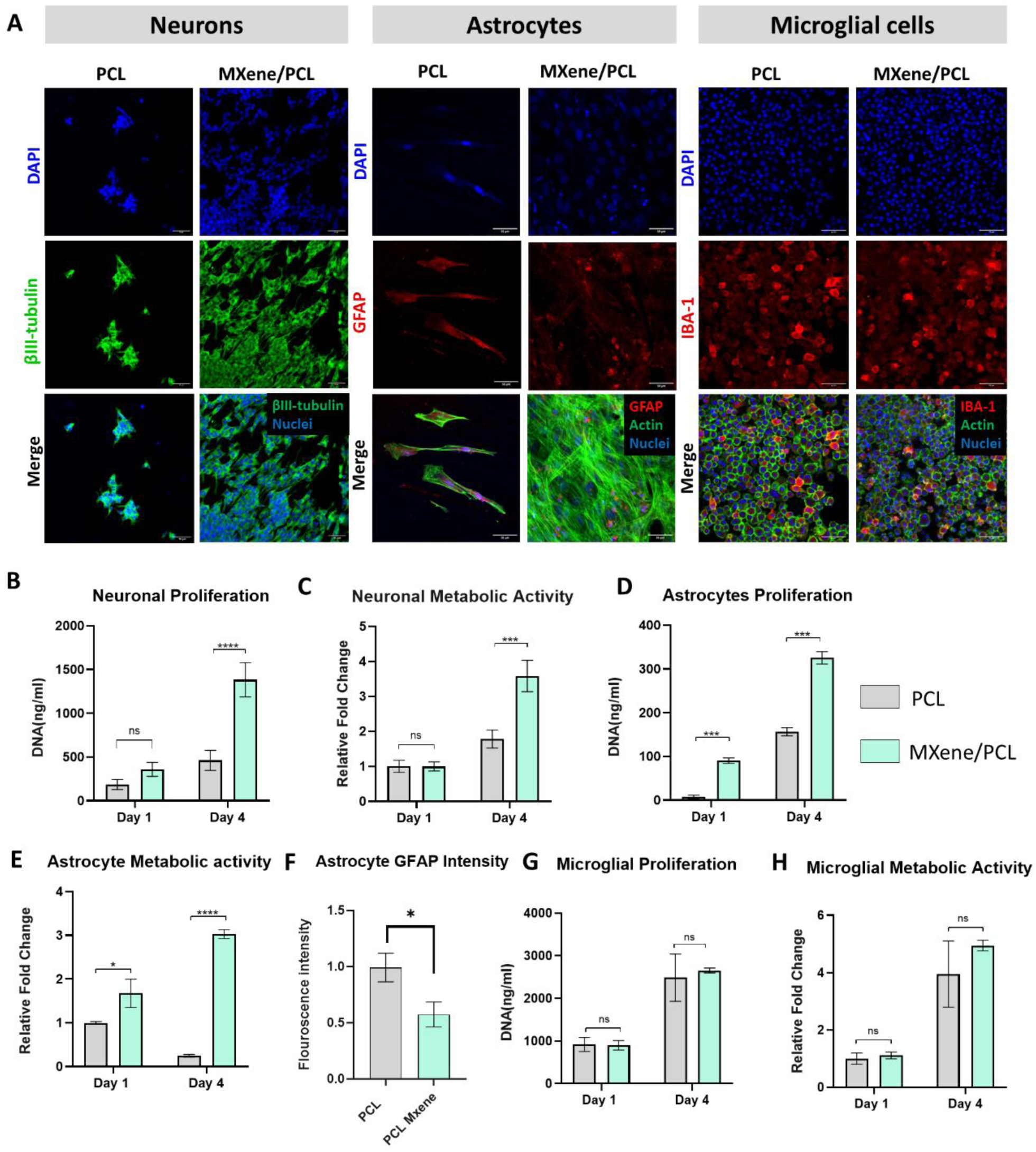
MXene/PCL Composite Biomaterial Interface Biocompatibility. (A) Immunohistochemical analysis of neuronal, astrocyte and microglial morphology when grown on the MXene/PCL or PCL-only substrates. (B) Neurons exhibit significantly increased proliferation on MXene/PCL surfaces and (C) increased metabolic activity. (D) Astrocytes exhibited poor attachment to the PCL surfaces but similar levels of proliferation and increased metabolic activity on both substrates (F) while also showing significantly (p<0.05) decreased GFAP expression on MXene surfaces. (G) Microglial exhibit no significant change in cell number or metabolic activity on MXene substrates. (Scale bar = 50 μm). *,**,*** indicates p <0.05, 0.01 and 0.001, respectively. p values calculated using Two-Way ANOVA with Tukey post-hoc testing, error bars indicate S.E.M.

### 2.3 Manufacture and characterization of MXene-functionalized MEW Micro-Meshes

To develop 3D MXene networks for incorporation within the tissue engineering scaffolds, melt-electrowriting (MEW) was used to 3D-print PCL microfibres with microscale resolution. The PCL micro-meshes acted as a template for functionalization with MXene nanosheets to form an interconnected conductive nanosheet-polymer network with a scale matched to the microanatomical features of native nervous tissue. For example, the corticospinal tract of the spinal cord exhibits maximum Feret diameters in the range of 0.5 – 1.5 mm (depending on location in the cord) while the tibial fascicle of the sciatic nerve exhibits a narrower diameter of approximately 500 μm^[19,45,46]^. To accommodate 3D-printing on this scale, melt electrowriting (MEW) was used. MEW provides significant advantages in terms of spatial control and fibre diameter over traditional fused deposition modelling approaches due to its microscale resolution and capacity to accurately print microfibrous polymer micro-meshes^[47]^. Shaper™ software (RegenHu, Switzerland) (Figure 3a) was used to design rectilinear mesh patterns with varying densities/spacings of microfibres, labelled as low, med and high density and representative of 1000, 750 and 500 μm spacings between fibres in each layer (Figure 3b). This rectilinear mesh architecture, beyond its scaling to approximate the size of anatomical features, also provides an isotropic structure which approximates the transverse parallel patterning of axonal tracts and bundles in the central and peripheral nervous systems^[19,45,46,48]^. Changing the fibre density resulted in significant changes in the effective cross-sectional areas of the scaffolds across each horizontal axis (Figure 3c), providing a mechanism through which to control the bulk electroconductive properties of the MXene micro-meshes. The direction of the infill was rotated by 90° for every second layer (Figure 3d) and the MEW parameters (Figure 3e) were optimized to produce high fibre resolution and an average fibre diameter (n=8) of approximately 13.64 ± 0.64 μm. For each 20 mm x 20 mm x 750 μm MEW print, several 6 mm diameter micro-meshes were punched out, which exhibited a high degree of spatial resolution (Figure 3f).

**Figure 3.**
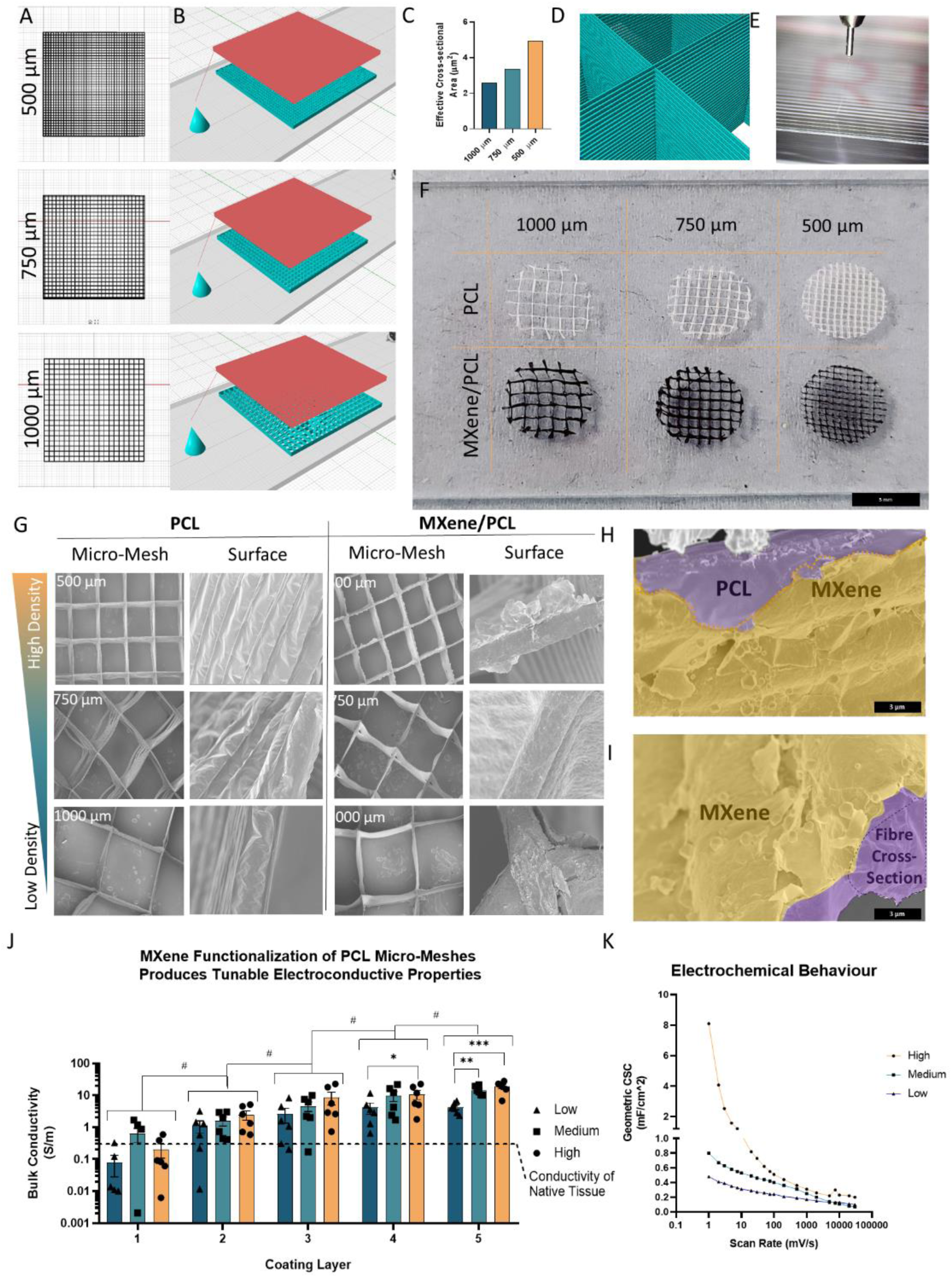
Melt Electrowriting of PCL/MXene Micro-Meshes. (A) Design of microfibre meshes (micro-meshes) of varying pore size. (B) SHAPR modelling of spinneret movement (red) and projected micro-mesh design (teal). (C) Variation in fibre spacing results in significant changes in micro-mesh cross-sectional area. (D) Micro-meshes are formed from interwoven layers of microfibers to a height of 75 layers. (E) Optimization of fibre deposition parameters results in high resolution printing. (F) Photos of 500, 750 and 1000 um micro-meshes. (G) Corresponding SEM micrographs of PCL and PCL/MXene micro-meshes, demonstrating effective production of 500, 750 and 1000 um microfibre-spacings. (H) MXene coatings result in homogenous adherence of MXene flakes to the PCL surface, which form (I) a thin intercalated crust. (J) The conductivity of MXene micro-meshes is dependent on microfiber density (p<0.001) and MXene layer thickness (p<0.001). (K) Analysis of the electrochemical behaviour of MXene microfiber architectures using cyclic voltammetry. *,**,*** indicates p <0.05, 0.01 and 0.001, respectively. # denotes significant p<0.05 effect of coating application on conductivity. P values calculated using Two-Way ANOVA with Tukey post-hoc testing, error bars indicate S.E.M.

Dip-coating of NAOH-treated PCL micro-meshes in 1 mg/ml MXene inks formed a black MXene film along the fibre surface. Subsequent SEM analysis of the PCL and MXene/PCL fibre architectures (Figure 3g) indicated that MXene coatings were spread evenly across the PCL fibres at each fibre density with the 2D flakes forming a thin scale-like crust across the fibre surface (Figure h, i). MXene nanosheets exhibited flat, flake-like morphologies, adhering in dense overlapping layers close to the fibre surface. Examination of the cross-sections of MXene-coated fibres demonstrated that the MXene treatment produced films consisting of multiple overlapping layers of MXene sheets. This layered film increased in consistency and thickness after each MXene ink application, resulting in significant increases in bulk conductivity across the scaffold (Figure 3j) with each coating (p<0.05). The conductivity of the low density micro-meshes increased from approximately 0.081 ± 0.053 S/m to 4.25 ± 0.67 S/m, while the medium density samples increased from 0.63 ± 0.3 S/m to 14.63 ± 1.93 S/m and the high density samples increased from 0.2 ± 0.1 to 18.87 ± 2.94 S/m, resulting in the highest bulk conductivity over-all. Statistical analysis indicated that both fibre density and coating layers significantly influenced the bulk conductivity of the MXene micro-meshes, indicating that their electrical properties were dependent both on the layering of MXene coatings and micro-mesh design, with conductivity roughly proportional to the cross-sectional area of fibres in the direction of measurement (Figure 3c).

Finally, the electrochemical properties of the scaffolds were analysed through cyclic voltammetry (Figure 3k) and measurement of the specific capacitance indicated that, despite the large volume of space within the MXene micro-meshes, their specific capacitance (0.3 – 1.05 mFcm^-^², low – high density), was equivalent to commercial platinum (approx. 0.1 – 0.2 mFcm^-^²)^[49,50]^, gold (0.11 mFcm^-^²) and CNT-based (1.51 mFcm^-^²)^[51]^ electrodes at 10 mV/s scan rates although significantly lower than carbon-fibre composites (e.g. PEDOT carbon fabric 70 mF/cm²)^[50]^. Despite this, the tunability of the system and the potential for higher density designs to be developed indicates that these MXene micro-meshes have significant potential for use in neural electrode applications.

### 2.4 Manufacture and characterization of biomimetic extracellular matrix-based (MXene-ECM) composite scaffolds containing MXene Micro-Meshes

MXene micro-meshes were then incorporated with a previously developed macroporous hyaluronic acid (HA)-containing ECM-based scaffold, functionalised with cord-native proteins, collagen type-IV (Coll-IV) and fibronectin (Fn). HA is the main structural component of the extracellular matrix of the central nervous system^[52]^ and previous work has shown that Coll-IV/Fn exhibit synergistic neurotrophic and immunomodulatory properties^[38,39,53]^. To produce these scaffolds, a 3mg/ml hydrazide-modified HA hydrogel was triturated with 100 μg/ml each of Coll-IV and Fn (Figure 4a), forming an ECM slurry^[39]^. PCL and MXene/PCL micro-meshes were then placed in 6 mm Teflon moulds with stainless steel bases and the micro-mesh filled with the ECM slurry. The metallic mould bases were flash-frozen to induce orientated ice crystal growth and then freeze-dried to form a macroporous ECM phase within the micro-mesh structures (Figure 4b). SEM analysis of PCL-ECM (Figure 4c) and MXene-ECM scaffold (Figure 4d) samples demonstrated the incorporation of the macroporous matrix throughout the micro-mesh structures.

**Figure 4.**
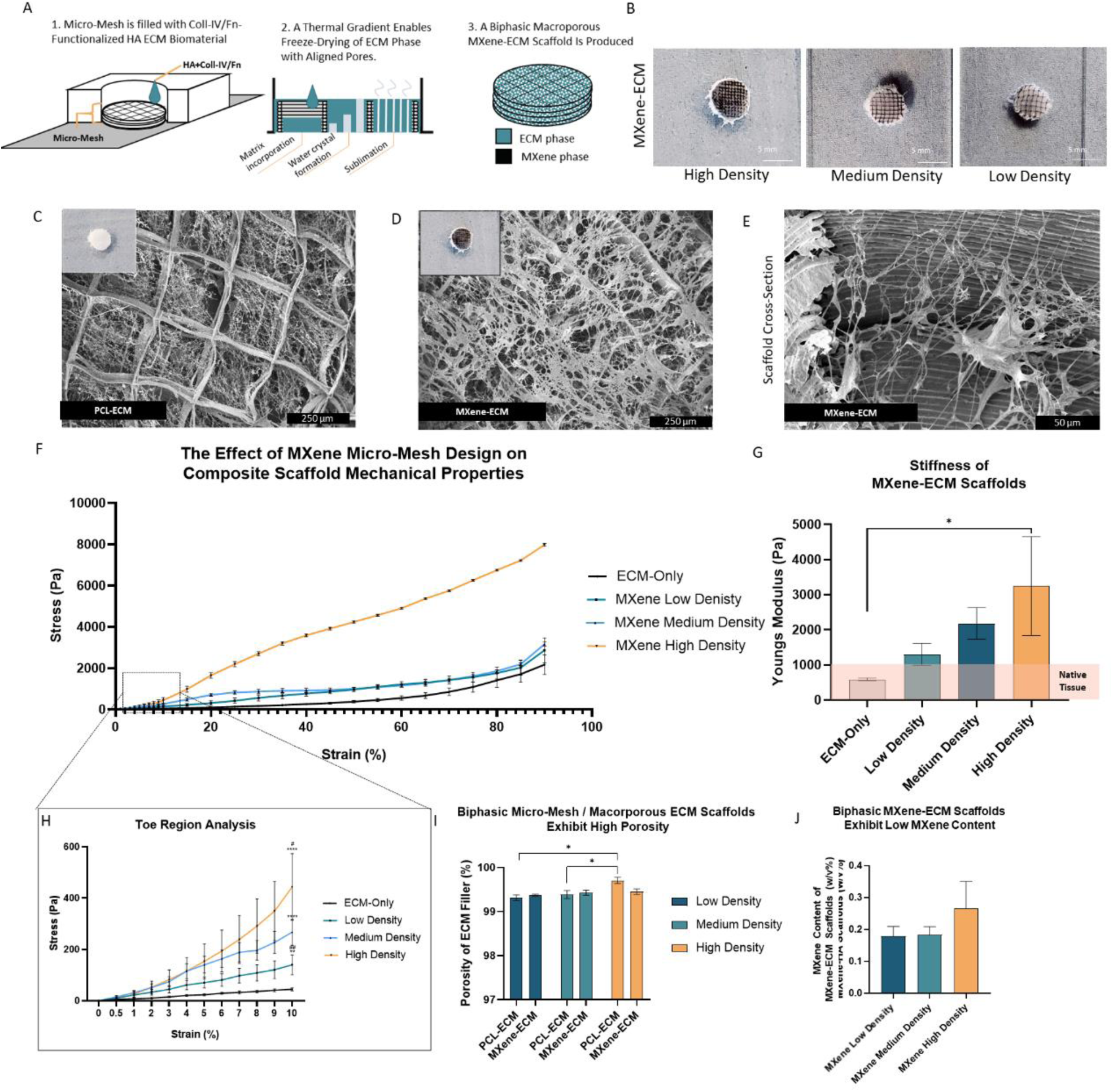
Freeze Drying and Characterization of HA-based ECM Within MXene Micro-Meshes to Produce Biphasic MXene-ECM Composite Scaffolds. (A) Schematic diagram detailing manufacturing process. (B) 500, 750 and 1000 μm ECM-filled micro-meshes. (C, D) SEM micrograph of biphasic scaffolds and cross-section (E). (F) Stress-strain curves of 3 mg/ml HA scaffolds and 1000, 750 and 500 μm biphasic MXene-ECM scaffolds. (G) Youngs Modulus of MXene micro-meshes. (H) Analysis of the stress-strain behaviour of composite scaffolds in low strain regions. (I) Proportion of ECM filler to micro-mesh. (J) Weight:Volume ratio of MXene content in biphasic MXene-ECM scaffolds. *,**,*** indicates p <0.05, 0.01 and 0.001, respectively. p values calculated using One-Way or Two-Way ANOVA, as appropriate, with Tukey post-hoc testing, error bars indicate S.E.M.

Examination of the scaffold cross-section indicated the formation of an orientated pore architecture in the spaces between the microfibrous micro-meshes (Figure 4e). Analysis of the stress-strain behaviours of the ECM-only scaffolds (Figure 4f) and composite MXene-ECM scaffolds indicated that increasing densities of MXene microfibres significantly increased the bulk stiffness of the scaffolds. The macroporous ECM-only material exhibited a biomimetic Young’s Modulus of approximately 0.6 ± 0.04 kPa (Figure 4g), matching the approximate softness of native brain tissue^[54,55]^ and within a similar range to what we have previously shown to promote neurite growth^[38]^. Low density MXene-ECM scaffolds exhibited a similar average modulus of approximately 1.3 ± 0.31 kPa which climbed to 2.18 ± 0.5 kPa for medium density samples and 3.25 ± 1.41 kPa for high density MXene-ECM scaffolds. Examination of the toe-region characteristics (Figure 4h) of these materials indicated that all groups exhibited similar mechanical properties, likely due to initial deformation of the outer ECM matrix coating at low strains before recruitment of the microfibres within the micro-mesh structures at higher strains. The average porosity of the macroporous ECM phase was above 99% in each sample (Figure 4i), indicative of a highly porous environment and the MXene content of the MXene-ECM scaffolds remained relatively low (Figure 4j), exhibiting an average MXene: Scaffold volume ratio of between 0.18 ± 0.03 % (low density) – 0.27 ± 0.08 % (high density).

### 2.5 Electrical stimulation of cell-seeded MXene-ECM composite scaffolds

To assess the efficacy of the electroconductive scaffolds in delivering electrical stimulation to neurons, the most conductive scaffold group, high density MXene-ECM scaffolds, were compared to MXene-free PCL-ECM control over a 7-day culture and stimulation period. Specifically, SH-SY5Y neurons were seeded on MXene-ECM scaffolds (n=12 per group) and cultured in differentiation medium for 7 days (Unstimulated) or moved to an IonOptix bioreactor after 24 hrs and stimulated for the next 6 days (Stimulated) (Figure 5a). Stimulated scaffolds exhibited robust cellular viability in all groups (Figure 5b) and metabolic activity increased in all groups throughout the stimulation period and was highest in the MXene-ECM, non-stimulated scaffolds. Confocal microscopy was used to analyse the morphology of the neurons and quantify the extension of βIII-labelled axons (Figure 5c) and excellent coverage of each scaffold and longer, denser neurites visible in both stimulated and non-stimulated MXene-functionalized samples. No significant change in cell number was observed between all groups (Figure 5d), but the per-cell metabolic activity was significantly lower (p<0.01) in stimulated MXene-ECM scaffolds compares to stimulated PCL-ECM samples (Figure 5e), consistent with maturation of SH-SY5Y neuronal cells towards a non-proliferative neuronal phenotype^[56]^. Analysis of the longest axon in each sample scaffold (Figure 5f) indicated that the longest axons were observed in the electrically stimulated MXene-ECM samples (108.5 ±6.13 μm), in which axons were significantly (p<0.01) longer, on average, compared to non-stimulated PCL-ECM controls (74.3 ±5.78 μm) and non-stimulated MXene-ECM scaffolds (67.41 ±6.52 μm). These results indicate electrical stimulation promoted axonal growth but no significant increases in maximum axonal length or average cellular neurite length (Figure 5g) were observed due to the conductivity of the MXene micro-mesh, although some significant changes in metabolic activity were detected.

**Figure 5.**
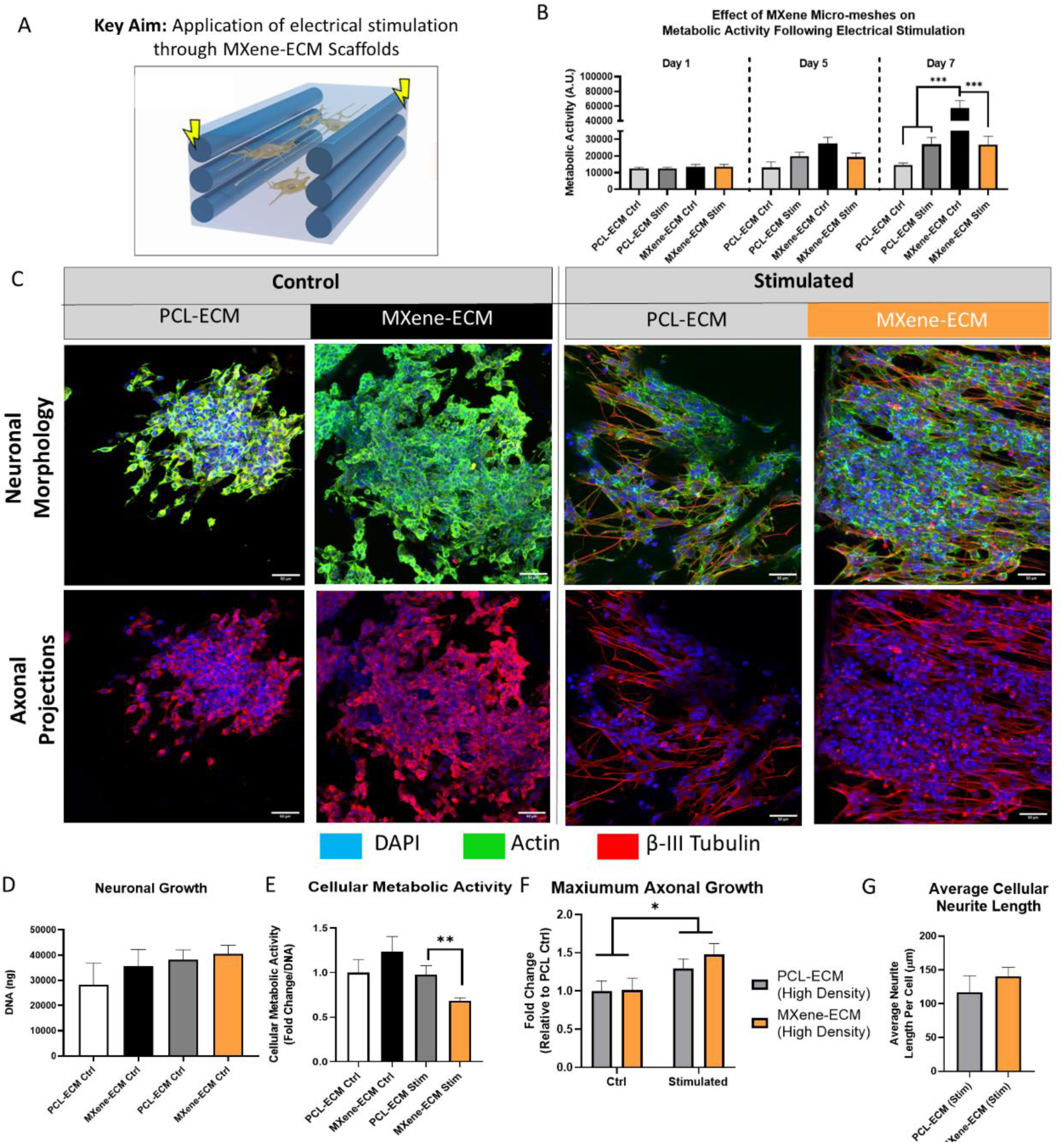
The Effect of MXene-ECM Biphasic Scaffold Conductivity and Micro-Mesh Design on Neuronal Cell Behaviour in Response to Continuous Electrical stimulation. (A) Schematic Diagram (B) Metabolic activity of neurons throughout long term electrical stimulation. (C) Immunohistochemical analysis of neuronal cell morphology (DAPI/Actin/BIII Tubulin). Neurons exhibit robust neurite projection in response to electrical stimulation. (D) DNA quantification following 7 days of culture and (E) corresponding metabolic activity at Day 7. (F) Electrical stimulation significantly enhanced axonal growth with High-density scaffolds exhibiting the longest axons. (G) Analysis of average neurite length per cell indicated no significant increase in neurons stimulated on MXene-HA scaffolds compared to inert PCL controls. Scale Bar = 50 μm. *, **, *** indicates p <0.05, 0.01 and 0.001, respectively. p values calculated using One-Way or Two-Way ANOVA, as appropriate, with Tukey post-hoc testing, error bars indicate S.E.M.

### 2.6 The influence of Micro-Mesh design on ES delivery to neurons in MXene-ECM composite scaffolds

Having examined how MXene micro-meshes effect delivery of electrical stimulation to neuronal cells, we subsequently examined how changes in the spatial organization of the MXene micro-mesh (high, medium and low density) within the scaffolds could enhance the axonal growth-promoting effects of the stimulation regime (Figure 6a). No significant difference between metabolic activity was observed between the different scaffold designs over time, although metabolic activity increased in all scaffold groups compared to Day 1 throughout the stimulation period, indicating that the stimulation regime was well tolerated by the cells (Figure 6b). Immunohistochemical analysis of neuronal morphology (Figure 6c) following 7 days of culture indicated the formation of extensive interconnected neurite networks in all samples, wherever dense cell numbers were observed. The DNA content (Figure 6d) in the stimulated scaffolds was proportional to the MXene/PCL fibre density, with high density samples exhibiting significantly higher DNA content compared to low density scaffolds. The highest average neurite length per cell (Figure 6e) was observed in medium density groups (365 ± 58.8 µm) compared to low (120.4 ± 12.1 µm) and high density (140.6 ± 12.93 µm) samples. These medium density scaffolds also exhibited significantly enhanced (p<0.05) βIII-tubulin expression compared to low and high density designs (Figure 6f), indicative of the increased maturation of the cell line neurons in these scaffolds^[56]^. Finally, analysis of the effect of MXene micro-mesh design on the maximal axonal length of stimulated cells (Figure 6g) indicated that higher density micro-meshes improved the axonal growth promoting characteristics when stimulated with medium density scaffolds exhibiting significantly longer (p<0.01) maximum axonal length compared to low density scaffolds although no significant differences in maximum axonal length between medium and high density samples were observed.

**Figure 6.**
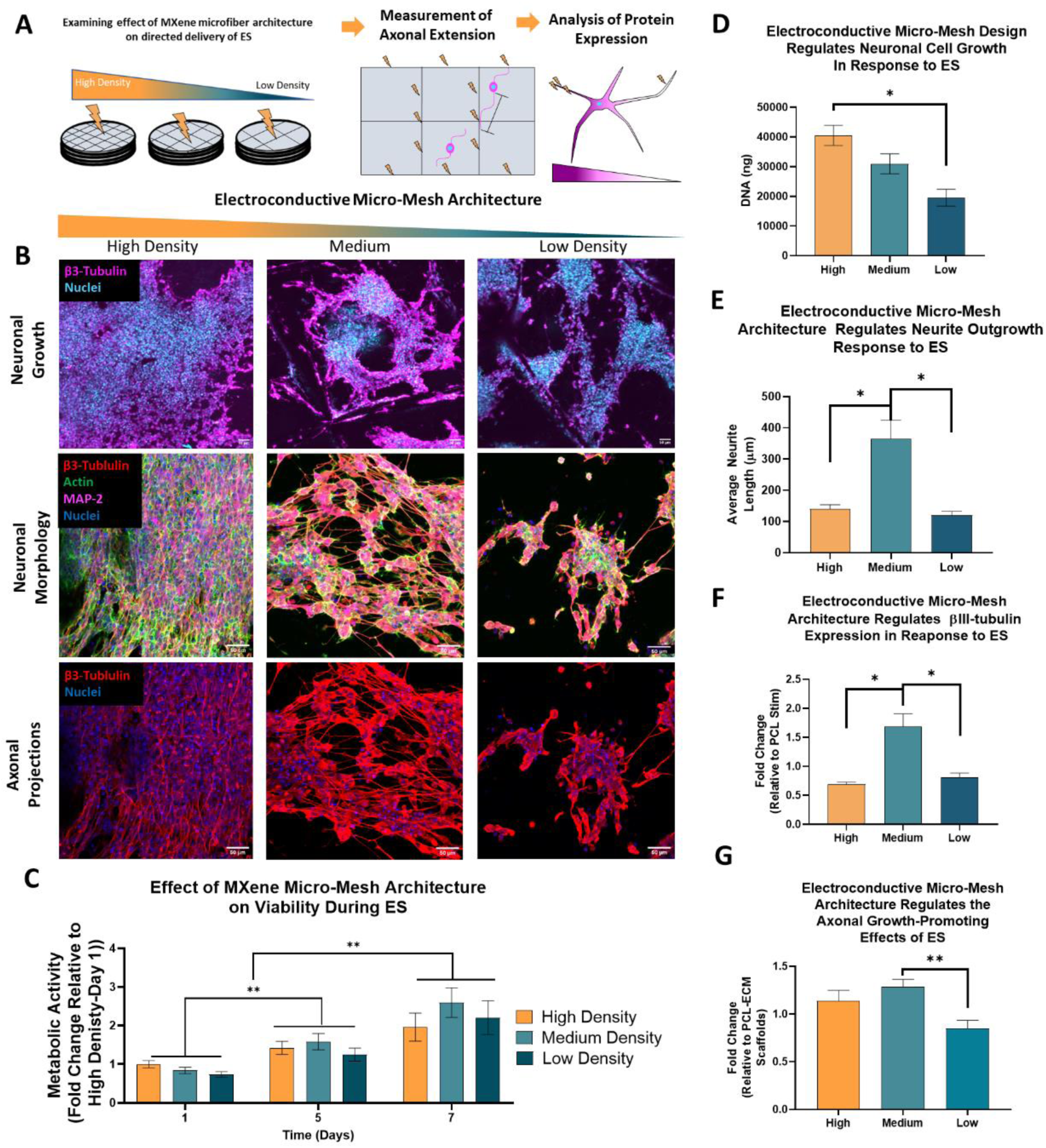
The Effect of MXene-ECM Scaffold Conductivity and 3D-printed Micro-Mesh Design on Neuronal Cell Behaviour in Response to Continuous Electrical stimulation. (A) Schematic Diagram of the Experimental Approach. Neuronal cells are seeded on MXene-ECM scaffolds, and following one week of culture with stimulation, imaged using immunohistochemistry to analyse both cell morphology and protein expression. (B) Immunohistochemical analysis of neuronal coverage, morphology and axonal extension. (C) Neurons on all MXene-ECM scaffolds exhibit increased metabolic activity throughout the stimulation period. (D) Cells exhibited significantly improved metabolic activity on High density MXene-ECM scaffolds. (E) Medium density scaffolds significantly enhanced the average neurite length per cell compared to Low and Medium density micro-meshes. (F) βIII-tubulin expression was significantly enhanced in Medium density scaffolds. (G) Maximum axonal length was significantly enhanced (p<0.01) in Medium density relative to Low density scaffolds. Scale bar = 50 µm. *, **, *** indicates p <0.05, 0.01 and 0.001, respectively. p values calculated using One-Way or Two-Way ANOVA, as appropriate, with Tukey post-hoc testing, error bars indicate S.E.M.

### 2.7 Electrical stimulation of olfactory bulb-derived stem cell neurospheres on MXene-ECM composite scaffolds

Having determined that micro-mesh design played a key role in the electroactive signalling capacity of the biphasic MXene-ECM scaffolds, we next sought to study the delivery of electrical stimulation in a more complex multicellular system through the use of neurospheres. The olfactory bulb represents a neuronal stem cell niche from which multipotent adult neural stem cells can be isolated which have the capacity to differentiate into both neuronal and glial cells and which have previous been trialled for human use in spinal cord injury repair^[57–59]^. Mouse olfactory bulbs were isolated from P0 – P3 mouse pups (Figure 7a), disassociated into single cell suspensions and allowed to form stem cell neurospheres over 10 days of culture. These neurospheres exhibited markers of stemness, Nestin and SOX-2 but also exhibited expression of neuronal marker βIII-tubulin and glial marker GFAP – indicative of multipotency (Figure 7b). These multipotent ONSC neurospheres were seeded on low, medium and high density MXene-ECM scaffolds and electrically stimulated for 7 days using the previously described stimulation parameters (Figure 7c).

**Figure 7.**
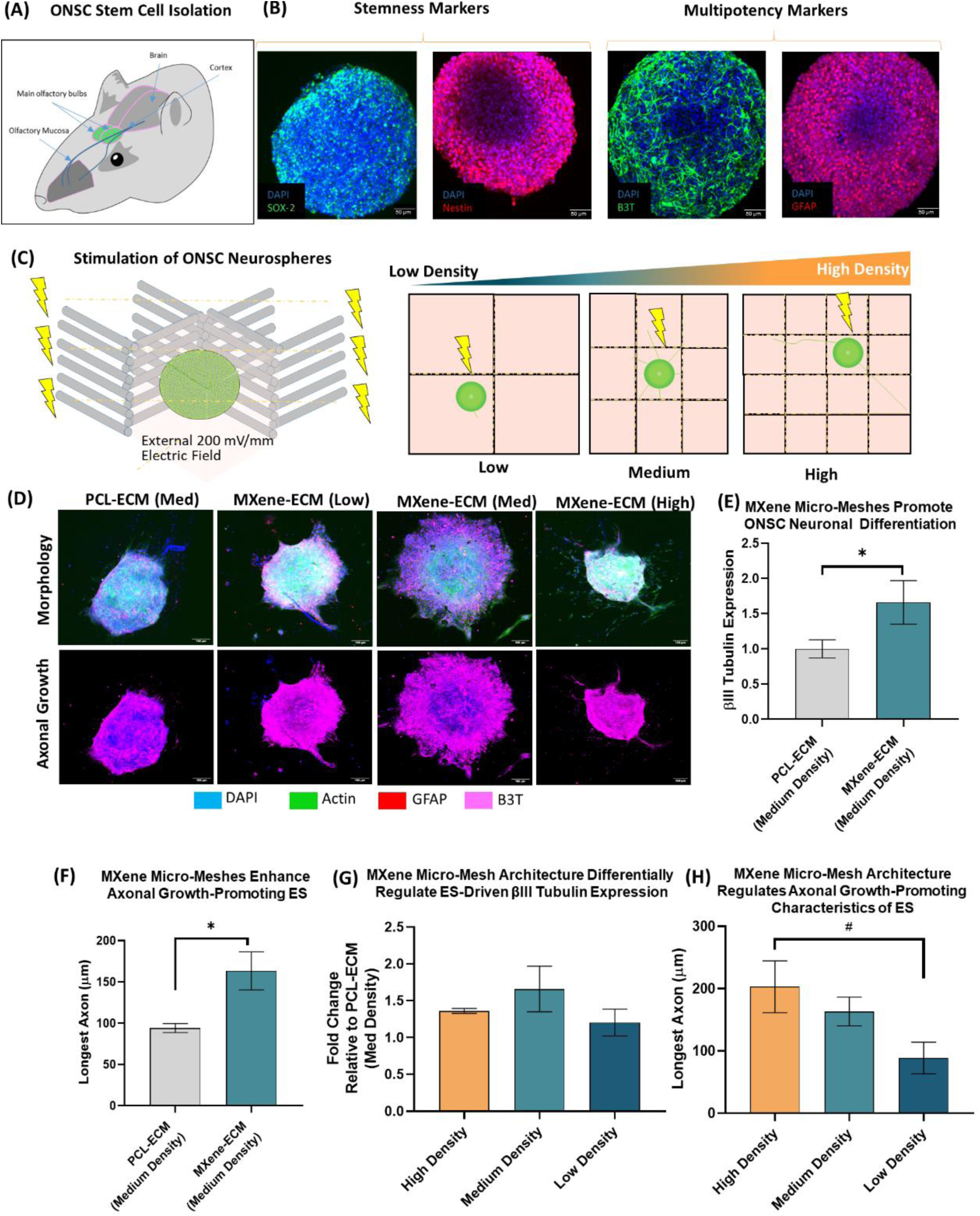
MXene-ECM scaffolds enhance the delivery of electrical stimulation to ONSC neurospheres in a manner dependent on 3D-printed Micro-Mesh design. (A) The olfactory bulb provides a source of adult olfactory bulb-derived neuronal stem cells (ONSCs). The culture of primary cells isolated from the olfactory bulb in suspension culture allows the isolation of stem cells in the form of neurospheres. (B) Confocal microscopic analysis of ONSC neurospheres indicates positive expression of positive markers of stemness (SOX-2 and Nestin) as well as markers of neuronal and glial differentiation (βIII-tubulin and GFAP, respectively) indicative of multi-potency (Scale bar = 50 µm). (C) Schematic diagram of neurosphere stimulation experiments. Individual ONSC neurospheres were seeded on High, Medium and Low density MXene-ECM scaffolds (n>4) and stimulated for 7 days. Immunohistochemical analysis was used to analyse axonal growth from the spheres and to quantify protein expression (D) ONSC neurospheres exhibit outgrowth of βIII-tubulin positive axonal processes across the scaffolds in response to electrical stimulation with increased outgrowth observed on MXene functionalized scaffolds (Scale Bar = 100 μm). (E) βIII-tubulin expression is increased on MXene-ECM scaffolds compared to inert PCL-ECM controls and (G) this trend is observed across all conductive microfiber designs. (F) The functionalization of microfibers with MXene significantly enhances the axonal-growth promoting effects of ES in Medium MXene-ECM scaffolds compared to PCL-ECM controls. (H) Increasing the density of MXene-microfibers significantly (p<0.05) enhances the neurite-growth promoting effects of the MXene-ECM scaffolds. Both Medium and High density MXene-ECM scaffolds drive the outgrowth of longer and thicker neuronal processes compared to Low density scaffolds. Annotations of *, **, *** indicates p <0.05, 0.01 and 0.001, respectively. p values calculated using Student’s t-test or One-Way ANOVA with Tukey post-hoc testing, error bars indicate S.E.M.

Immunohistochemical labelling and confocal microscopy were used to analyse and cellular differentiation and the outgrowth of axons from the neurosphere (Figure 7d). The sphere’s exhibited increased outgrowth on the medium density MXene-ECM scaffolds compared to those stimulated on non-electroconductive PCL-ECM controls with the projection of long axons in all directions and the outgrowth of cells into the surrounding biomimetic matrix. The delivery of electrical stimulation to spheres on the MXene-ECM scaffolds (medium density) enhanced βIII-tubulin expression (Figure 7e) by approximately 2-fold (p<0.05) and enhanced max axonal length (Figure 7f) increasing from 94.14 ± 5.47 µm (PCL-ECM, medium density) to 163.4 ± 22.98 µm (MXene-ECM, p<0.05). Furthermore, when the effect of stimulation on key therapeutically-relevant behaviours was analysed, the results demonstrated that the electroconductive micro-meshes within the MXene-ECM scaffolds played a significant role in the delivery of electrical stimulation. Neuronal differentiation was broadly improved across all MXene-ECM scaffolds compared to the inert PCL-ECM ctrl group, although no significant difference between groups was observed (Figure 7g). However, the higher density MXene-ECM scaffolds exhibited the longest axonal extension (Figure 7h) 203 ± 41.51 µm, significantly (p<0.05) higher than both PCL-ECM controls (94.14 ± 5.4 μm) as well as the low density MXene-ECM scaffolds (88.6 ± 25.5 µm). These results demonstrate that the spatial organization of the MXene micro-mesh within the biphasic MXene-ECM scaffolds had a significant effect on the axonal growth-promoting aspects of the electrical stimulation and this effect was observed across both cell line neurons and primary neuronal stem cells.

## 3. Discussion

Neurotrauma injuries such as spinal cord or traumatic brain injury present a complex challenge for repair and no treatment currently exists that can effectively repair damaged CNS tissues. In this study we have successfully developed a multifunctional tissue engineered scaffold whose tunable electroconductive properties can be adapted through microscale 3D-printing to tailor and thus enhance the pro-reparative effects (including increased axonal length and density, as well as enhanced neuronal differentiation) of therapeutic electrical stimulation for neural repair applications. Incorporating the MXene/PCL micro-meshes within a biomimetic HA-based, Coll-IV/Fn-functionalized scaffold produced a multifunctional biphasic composite MXene-ECM scaffold. This scaffold exhibited soft mechanical properties, high electroconductivity (despite a low overall MXene content) and a bioactive macroporous phase that facilitates biological functionality and axonal growth in response to electrical stimulation. The application of stimulation promoted axonal growth from seeded neuronal and stem cells in a manner dependent on the electroconductive scaffold micro-mesh architecture and demonstrates that 3D-printing can be utilized to significantly enhance the electroactive characteristics of tissue engineering scaffolds.

In order to establish whether microscale patterning of conductive biomaterials through 3D-printing could be used to direct the delivery of electrical stimulation to neurons and stem cells in a tissue engineering scaffold, a sequential design process was utilised. Ti_3_C_2_T*_x_* MXene nanosheets were initially derived from an aluminium (Al)-rich MAX phase precursor which provides degradation resistant particles with excellent electroconductivity and robust hydrolytic stability^[41]^. The resultant MXene films exhibited excellent conductivity (>10,000 S/cm) – indicative of the high electroconductivity properties of the MXenes. These values were several orders of magnitude higher than previous studies that have used biocompatible MXenes for cellular electrical stimulation experiments^[17,37]^, with the observed increases likely due to differences in the composition of the MAX phase precursor material, the size range of the prepared flakes^[60]^ and the methodology for film preparation (vacuum annealing compared to spin-coating). Ti_3_C_2_T*_x_* MXene materials have been shown to provide excellent biocompatibility in a range of cell culture applications^[40]^. For example, MXene films have previously been used to effectively deliver ES to iPSC-derived cardiomyocytes^[37]^ and neuronal progenitor cells to enhance their maturation^[17]^. Furthermore, work by Zhu et al. (2023) demonstrated that subcutaneously implanted MXene-coated electrospun PLLA microfibres do not invoke a strong foreign body response^[61]^. However, the interaction of MXenes and MXene/PCL composites with glial cell types that play important roles in neural inflammation has not been widely investigated^[40]^. This study shows that MXene/PCL surfaces possess excellent biocompatibility with neurons, astrocytes and microglia, indicative of the suitability of the MXene coatings for implantation in neural tissue. Furthermore, the observed decreases in GFAP expression in astrocytes indicated that the MXene surface did not promote the adoption of a reactive astrocyte phenotype which is a key challenge for the compatibility of biomaterial interfaces with neural tissues^[62]^.

To achieve the necessary spatially-controlled distribution of the conductive MXene/PCL materials throughout the scaffold, melt electrowriting was used to 3D-print highly organized rectilinear microfibrous PCL micro-meshes with varying fibre spacings producing scaffolds of low, medium and high fibre density (corresponding to 1000, 750 and 500 µm spacings, respectively). MEW provides a combination of 3D-printing and electrospinning that allows the “direct writing” of polymer melt jets to produce 3D microfibrous micro-meshes^[47]^. Functionalization of these micro-meshes with MXenes resulted in a highly contiguous network of intercalated MXene flakes coating the fibre surfaces, resulting in a bulk conductivity in the high density samples far higher (approx. 19 S/m) than native tissue (approx. 0.4 S/m^[63]^). By confining the MXene nanosheets within the scaffold to dense intercalated coatings of the slow-degrading PCL microfibre micro-meshes^[64]^, the volume ratio of MXenes required to produce a conductive network throughout a scaffold was efficiently minimized (<0.5 w/V%). Additionally, tuning the geometry of the MXene micro-meshes allowed the control of their electrochemical properties - achieving relatively high geometric capacitance in the high density design comparable to widely used electrode materials^[65]^, despite their high porosity and flexible mechanical properties demonstrating the suitability of MXene/PCL fibres as a promising material for flexible electrode design.

MXene-PCL scaffolds have been previously developed to deliver optocapacitance-based neural stimulation^[66]^, whereas conductive MEW scaffolds have been manufactured utilizing conductive polymers (e.g., polypyrrole)^[67]^ and metallic (e.g., gold)^[68]^ coatings. However, a key limitation of the use of conductive polymers (such as polypyrrole) is the rapid degradation due to oxidation^[69]^ and leaching of dopants, while metals exhibit high stiffness – leading to scarring. In contrast, the Al-rich MAX-phase-derived Ti_3_C_2_T*_x_* MXenes utilized in this study exhibited robust degradation properties. Degradation testing of the MXene films and MXene-ECM scaffolds in PBS solutions, designed to mimic the oxidative and hydrolytic processes that occur *in vivo*, showed that the films and scaffolds exhibited the retention of approximately 50% of their electroconductive properties after 7 days. The loss of conductivity is likely to be due to swelling and physical disruption of the mechanically-weak hydrophilic films, resulting in disruption of contacts between the MXene nanosheets, rather than chemical degradation of the MXene nanosheets through hydrolysis or oxidation. However, even considering these decreases, the freestanding MXene films retain excellent electroconductivity and these results indicate significant potential for the use of high density MXene-microfibre biomaterials in neural electrode applications e.g., electrocorticography recording devices^[49,55,70]^.

The incorporation of the MXene/PCL micro-meshes within the macroporous biomimetic HA-based matrix provides a highly neurotrophic biphasic (MXene-ECM) scaffold which combines the spatial control of melt electrowriting and the tunable electroconductive qualities of the MXene composite fibres with the neurotrophic and immunomodulatory properties of our previously developed HA-Coll-IV/Fn biomaterial^[38,53]^.This biomimetic HA Coll-IV/Fn macroporous filler material exhibits a similar softness to the native brain tissue^[71]^ and the functionalization of HA with collagen type-IV and fibronectin synergistically enhances axonal growth – making it an excellent substrate for trialling therapeutic electrical stimulation strategies^[38]^. Melt electrowritten microfibre micro-meshes are most commonly used to integrate physical cues within tissue engineering scaffolds^[42,72,73]^ or to provide mechanical reinforcement of soft materials^[66]^. For example, previous studies have integrated MEW frameworks within Matrigel or thiolated HA hydrogels to produce soft but robust biomimetic substrates for modelling CNS tissue behaviours^[74–76]^. Advancing on these approaches, this study utilizes MXene/PCL micro-meshes to direct electrical stimulation rather than mechanical stress throughout the biphasic scaffolds while the soft biomimetic HA scaffolds provide the requisite soft mechanical properties in conjunction with bioactive signalling from the neurotrophic ECM to enhance pro-reparative cell behaviours. Even in high density samples with relatively high proportions of MXene/PCL microfibres, the bulk compressive modulus was close to the biomimetic range (approx. 3 kPa) while low density scaffolds exhibited an approximately biomimetic modulus in a similar range as native CNS tissues (0.5-3 kPa) ^[77–79]^.

In the biphasic MXene-ECM composite scaffolds, the majority of neurons were not in direct contact with the conductive material phase, nevertheless, the results show that MXene micro-meshes broadly influenced ES delivery to neurons throughout the scaffold. Previous work by Zhang et al.(2018) has shown that patterning of 3D electrodes in 3D biomaterial environments can manipulate electrical fields to influence axonal growth and cell aggregation^[9]^. In recent work, 3D-printed scaffolds containing polypyrrole-PCL cylindrical arrays, designed to mimic the anatomical geometries of spinal cord axonal tracts, effectively delivered ES to neurons in a macroporous HA filler material^[19]^. The present study not only demonstrates that the electroconductive MXene-ECM scaffold can enhance delivery of electrical stimulation to neurons but also demonstrates that the capacity of 3D-printing to control the spatial distribution of the electroconductive MXene phase within the biphasic composite could further enhance the electroactive signalling potential of these tissue engineering scaffolds - providing a novel role for 3D-printing in biomaterial design. Furthermore, this biphasic approach overcomes several challenges associated with the biofabrication of conductive biomaterials to produce soft, biomimetic highly conductive scaffolds with robust degradation properties.

The electrical stimulation of the SH-SY5Y neuronal cells was shown to significantly increase maximum axonal length in both the inert and MXene-functionalized high density scaffolds, demonstrating the benefits of applying the described ES regime (200 mV/mm, 12 Hz, 50 ms pulse). While this stimulation regime has previously been shown to promote axonal growth^[15,19]^, the stimulation regime applied through the MXene micro-meshes could be adjusted to target other cell behaviours such as supporting cell signalling or promoting neural plasticity^[80]^. Neurons stimulated on medium density MXene-ECM scaffolds exhibited a significant 3-fold increase in average neurite outgrowth per cell and significantly enhanced βIII-tubulin expression compared to high and low density design MXene-ECM scaffolds and maximum axonal length was significantly increased relative to low density scaffolds. These results agree with a recent study by Das et al (2024) which demonstrated that increasing the graphene content of electrospun silk nanofibre membranes could result in a loss of axonal growth promoting responses to electrical stimulation in seeded neurons^[81]^. Together these results suggest that neurons are highly sensitive to small changes in the electroconductive properties and spatial organization of the surrounding microenvironment and demonstrates that the relationship between micro-mesh design, electroconductive properties and the applied ES regime requires optimization beyond maximizing the electroconductivity of the scaffold.

To further investigate how tuning the electroconductive micro-mesh design influences the delivery of electrical stimulation to neural cells, the application of ES to ONSC-derived neurospheres was examined. ONSCs represent a therapeutically-relevant and potentially autologous source of neural stem cells with the capacity to adopt neuronal and glial subtypes, which have previously undergone human trials as a treatment for SCI^[57]^. Neurospheres enable *in vitro* testing to be carried out in conditions that more accurately model the cell diversity, complex-cell-cell interactions and 3D behaviours that occur in neural tissues^[82]^. These spheres were demonstrated to contain multipotent neural progenitor cells, capable of differentiating into neuronal and glial phenotypes, and exhibited excellent compatibility with the neurotrophic ECM scaffold phase. Stimulation of the ONSC-derived neurospheres promoted increased neuronal differentiation and axonal growth in MXene-functionalized medium density scaffolds compared to inert controls and this effect was further increased in high density scaffolds and decreased in low density scaffolds – demonstrating the significant effect of electroconductive micro-mesh design in altering the response of the neurospheres to an applied ES, although this trend differed from the improved response of neuronal cells on medium density micro-meshes. The response of cells to electrical stimulation is known to be cell type-specific^[80]^, and these results may speak to both the different environment cells experience within a multicellular neurosphere^[83]^ as well as a need to optimize micro-mesh design for cell-specific behaviours or tissue-specific applications.

In summary, this study shows that microscale control of the electroconductive properties of a suitable tissue engineering scaffold can have a significant effect on repair-relevant neuronal cell responses and that the combination of 3D-printing and effective use of nanocomposite materials can provide a highly effective means of enhancing the delivery of electrical stimulation to neuronal cells. While the synergistic interactions between conductive micro-meshes and ES to promote axonal growth observed in this study have important implications for conductive implant design, it is possible that the stimulation profile could be optimized for other aspects of neural regeneration, for example targeting electroactive cell types such as astrocytes or oligodendrocytes to modulate inflammation^[84]^ or promote re-myelination^[85]^, or even for the regeneration of other tissue types – such as bone, tendon or cardiac tissues.

## 4. Conclusion

This study describes the development of a 3D-printed, multifunctional, composite scaffold containing an electroconductive MXene-functionalized micro-mesh, whose electroconductive properties are highly tunable, embedded within a neurotrophic ECM-based scaffold. By microscale adjustment of the architecture of these MXene micro-meshes using melt-electrowriting, their capacity to deliver electrical stimulation was significantly enhanced, improving the axonal growth-promoting and pro-neuronal differentiation signalling effects of the applied stimulation. The findings highlight that tuning the interplay between electroconductive biomaterial distribution, electroconductive properties, electrical stimulation parameters, and target cell type is crucial, requiring optimization beyond maximization of scaffold conductivity. These results demonstrate a novel use of the spatial control of 3D- printing approaches in tissue engineering scaffold design with significant implications for the future development of electrically-active neural implants for the repair of lesions associated with neurotrauma.

## Acknowledgements

This study was funded by a joint funding initiative of the Irish Rugby Football Union Charitable Trust (IRFU-CT) and the Research Ireland Advanced Materials and Bioengineering Research (AMBER) Centre (SFI/12/RC/2278) and an Irish Research Council Postdoctoral Fellowship (Government of Ireland), Grant Number: GOIPD/2021/262, an EPSRC/ Research Ireland Centre for Doctoral Training in the Advanced Characterisation of Materials (18/EPSRC-CDT/3581 15735). The authors thank the Advanced Microscopy Lab (AML) at the Centre for Research on Adaptive Nanostructures and Nanodevices (CRANN) for their help with the performance of electron microscopy as well as our Public Patient Involvement group for their guidance and support.

## 4. Methods and Materials

### 4.1 Materials

All reagents were purchased from Sigma Aldrich (Ireland) unless stated otherwise. Wash steps refer to 3× Dulbecco phosphate buffered saline (DPBS) washes for 5 min at room temperature (RT) unless stated otherwise. All culture conditions were at 37 °C, 5% CO2 unless stated otherwise.

### 4.2 MXene Synthesis and particle characterization

To a 200 ml HDPE bottle, 9M HCl (40 ml) was added. A PTFE stirrer was placed inside and set to stir at 300 rpm in an ice bath. Once cooled, around 10 minutes, Al-rich Ti_3_AlC_2_ MAX phase powder (4 g, Carbon-Ukraine ltd.) was added in small additions (c. 400 mg/min) to avoid excessive heating of the solution. The lid was placed loosely on the bottle and the mixture stirred for 18 h to remove excess metals (mostly Al) from the powder. The mixture was then washed several times with deionised water, centrifuging at 5000 rpm using a Thermo Scientific Heraeus Multifuge X1 for 5 minutes each time, until the pH of the supernatant was > 6. The washed MAX phase sediments were then dried via vacuum filtration overnight. The resulting MAX phase powder was then weighed, typically having lost around 25% of its original mass.

To a 200 ml HDPE bottle in a mineral oil bath, 9M HCl (60 ml), followed by LiF powder (4,8g, Sigma) was added, and left to stir using a PTFE stirrer bar for 5 min. The washed MAX phase powder (3g) was then added slowly over 5 min. The solution was then left stirring at 400 rpm, 35°C for 24 hours to obtain the etched, multilayer Ti_3_C_2_T*_x_* MXene.

To wash, the contents of the vessel were transferred evenly into 2 x 50 ml centrifuge tubes and diluted to a total of 40 ml each with deionised water. The dispersions were then washed around 10 times via centrifugation at 5000 rpm until the pH of the supernatant was approximately neutral. The tubes were again filled to 40 ml with deionised water and vortex mixed for 1 hour at 1800 rpm to delaminate the MXene. The MXene dispersion was then centrifuged at 1500 rpm for 30 minutes to sediment any multi-layer MXene or unreacted MAX phase that remained. The supernatant containing delaminated MXene flakes was then collected. This supernatant was then centrifuged at 5000 rpm for 1 hour to sediment the few-layer flakes, the sediments were then re-dispersed in a total of 25 ml deionised water to obtain a concentrated MXene ink.

### 4.3 PCL Film manufacture and MXene Ink functionalization of the films

PCL (50 kDa, Polysciences Inc. PA, USA) was dissolved at a concentration of 20 wt% in chloroform before being cast on a Teflon mould to form a thin film. The thin film was allowed to air dry overnight before being punched into 6 mm discs. Each film sample was then treated with 3M NAOH overnight to etch the film surface and improve hydrophilicity. The PCL films were then dip coated in a 1 mg/ml MXene ink and allowed to dry overnight (x5 coating layers) to produce a MXene-functionalized PCL (MXene/PCL) film. These films were then stored at RT in an airtight container.

### 4.4 Electron Microscopy

SEM analysis of the pore structure of freeze-dried scaffolds was carried out similarly to previously published analyses^[38]^. Scaffolds were bisected in transverse and longitudinal orientations using microtome blades and prepared for imaging through gold sputter coating. Samples were inserted into the chamber of a Zeiss Ultra Plus field-emission SEM electron microscope (Zeiss, Oberkochen, Germany) and imaged using accompanying SmartSEM software. EDX of the Ti_3_C_2_T*_x_* was performed at an acceleration voltage of 10 keV with a 20mm² Oxford Inca EDX detector.

### 4.5 Surface Analysis

#### 4.5.1 Hydrophilicity

The contact angle of each group was determined by a goniometer (optic contact angle measuring and contour analysis system model OCA 25). A water droplet of 100µl is placed upon each film and the contact angle measured.

#### 4.5.2 Atomic Force Microscopy

To assess the surface topography of the MXene/PCL composites, AFM was carried out on a scanning probe microscope (NanoSurf) in tapping mode under ambient conditions using aluminium-coated silicon cantilevers (TAP-190i). Average image sizes were 50 × 50 μm at scan rates of 0.5 Hz with 512 lines per image. Images were analysed using CoreAFM software (NanoSurf).

### 4.6 Melt Electrowriting and MXene-functionalization of PCL Micro-Meshes

A RegenHu R200 3D-printer with a melt electrowriting (MEW) module was used to produce the microfibre scaffolds. Scaffold micro-meshes were designed on Shaper CAD Software (RegenHu). The fibre design was a rectilinear pattern with varied fibre spacings of 500, 750 and 1000 μm. The orientation of the layers was swapped perpendicularly after each layer to produce a “box-like” architecture. A 50 kDA polycaprolactone polymer was used for microfibre manufacture. The following Mew parameters were used: 5 mm tip-to-collector distance, 10 mm/s translation speed, approximately 5 kV voltage, 50 kDA PCL, heated to 80 °C, 24 G spinneret and a 50 kPa extrusion pressure. The Taylor cone was allowed approximately 10 minutes to stabilize prior to each print. The patterns were printed to a 75- layer height with 10 μm diameter fibres to produce 750 μm-tall PCL micro-meshes. Fibre diameter (n=6) was measured using scanning electron microscopy.

To functionalize the PCL micro-meshes with a MXene coating, the micro-meshes were treated with 3M NAOH for 4 hrs before being washed with PBS and then deionized H_2_0. The treated scaffolds were then dip-coated in a 1mg/ml MXene ink, and allowed to dry, forming a MXene/PCL micro-mesh. This dip-coating process was repeated 5 times to increase the uniformity of the coating and the electroconductive properties of the micro-meshes.

### 4.7 Electrical Properties

#### 4.7.1 Conductivity Analysis

Film conductivity measurements were performed using an Ossila four-point probe, with a probe spacing of 1.27 mm. Bulk electrical conductivity was determined using the two-point probe method with a probe head (SP4-40180TRJ, Lucas Signatone) connected to a source meter instrument (Model 2450, Keithley). 20 mm x 20 mm microfibre micro-meshes were placed between two carbon electrodes and reading made in a dry state at RT. Current readings were obtained by sourcing voltage in the ±3 V interval to screen the linear region of the films according to Ohm’s law.

#### 4.7.2 Cyclic Voltammetry

Two-electrode coin cells were fabricated with 3 mm discs of electrode material. Cyclic voltammetry (CV) and electrochemical impedance spectroscopy (EIS) measurements were carried out using a Biologic VMP-300 potentiostat. CV was carried out in a potential window of ±200 mV at scan rates ranging from 1 mV/s - 30000 mV/s. EIS was carried out in the frequency range 1 Hz – 7 MHz under a potential amplitude of 10 mV. Charge storage capacity and charge transfer resistance analysis was carried out using Biologic EC-Lab software.

### 4.8 Freeze-drying of macroporous microfibre composite scaffolds

A Hyaluronic acid hydrogel was prepared using a modified carbodiimide chemistry process essentially as previously described^[38,39]^. Briefly, HA sodium salt (1.6–1.8 MDa, Streptococcus equi, 53747) was dissolved in dH20 at a concentration of 3 mg/ml. Adipic acid dihydrazide (ADH) was added as a crosslinker in excess (1.832 g ADH/ 0.5 g HA) while 1-ethyl-3-(3- dimethylaminopropyl) carbodiimide (EDAC) was used to activate the HA carboxylate groups (800 mg per 1 g HA). The reaction was initiated by dropping the pH to 4 using 1 m HCL and crosslinking was carried out overnight. The next day the pH was adjusted to pH 7 and the HA solution was dialyzed in dialysis tubing (6–8 kDa MWCO) through 100 × 10−3 m NaCl for 24 h (x2), followed by 20% ETOH and finally dH20. At this point, collagen type-IV (human placenta, C5533) and fibronectin (human plasma, FC010) solutions were triturated at a concentration of 0.1 mg mL−1 within the HA solution to form an extracellular matrix-based (ECM) biomaterial. Subsequently, microfibre frameworks were placed within Teflon moulds, filled with the ECM solutions and freeze-dried (TF =-40 °C). The macroporous composite scaffold was formed via the freezing of the ECM solution, phase separation of the water into crystals, compaction of the HA polymer chains and evaporation of water crystals to create a porous structure. Scaffolds were subsequently rehydrated through 100% and 70% ETOH solutions overnight and immersed in an EDAC/N-hydroxysuccinimide solution to crosslink the ECM material within the freeze-dried MXene/PCL and PCL micro-meshes. The resulting PCL- ECM or MXene-ECM scaffolds were washed with DPBS before being stored overnight at 4 °C prior to cellularization or characterization. Mechanical testing of the HA scaffolds was carried out in a Zwick mechanical testing unit in a uniaxial compression configuration in a PBS bath. Cylindrical HA samples were tested to 60% strain using a 5N load cell using a displacement rate of 100% strain/min and the Youngs Modulus calculated using the slope of the stress-strain curve between 1% and 10% strain.

### 4.9 Biocompatibility

Primary human spinal cord astrocytes (Isolated and cryopreserved by ScienceCell, USA) were grown in DMEM-Ham’s F12 supplemented with 1% PS, FBS, and astrocyte growth supplement (ScienceCell, USA). Following plating, cells were cultured in growth media for 1 day before changing the media to a serum-free formula of DMEM-Ham’s F12, 1% PS, L-Glutamine, N21 max (R&D Systems, UK), and 0.2% B27 supplement. To assess astrocyte compatibility, human astrocytes (P3-6) were seeded at a density of approximately 10,000 cells per well in 96 well plates containing 6 mm PCL or MXene/PCL films (n=4). Immortalized microglial cells (IMGs) were used to model the compatibility of the MXene surfaces with microglia cells, due to the role the microglia have in activating inflammation responses in the brain. IMGs were seeded at a density of 100,000 cells per well in 96 well plates containing 6 mm PCL or MXene/PCL films. IMG growth medium consisted of DMEM-Ham’s F12, 1% PS, 1% L-Glutamine and 5% FBS. SH-SY5Y neuronal cells (American type culture collection, USA) were used to model the compatibility of the MXene surfaces with neurons. 100,000 cells per well of P12-P16 neurons were seeded on 6 mm PCL or MXene/PCL films. Growth medium consisting of DMEM:Ham’s F12 (50:50) containing 10% fetal bovine serum (FBS, Biosera, Ireland), 1% L-glutamine, and 1% penicillin-streptomycin (PS). In each case, cell-seeded films were cultured for 1 - 4 days in growth medium, with Alamar Blue assay-based analysis of metabolic activity at days 1 & 4 before being washed with PBS and fixed for immunohistochemical analysis using 4% PFA or frozen for DNA analysis.

### 4.10 Electrical Stimulation of MXene-ECM macroporous scaffolds

MXene-ECM scaffolds were manufactured with 500, 750 or 1000 μm fibre spacings. PCL- ECM controls were manufactured with 500 μm spacings, only. SH-SY5Y cells were seeded at a density of 500,000 cells per scaffold in 6 mm PCL-ECM or MXene-ECM scaffolds filled with the ECM macroporous material. Following 24 hrs of culture in growth medium to enable cellular infiltration of the scaffolds, the cellularised scaffolds (n=6) were transferred to 8 well plates and stimulated using protocols previously developed for the promotion of axonal growth^[15]^. Using an in-plate electrode system (8-well plate C-Dish, IonOptix), a 200 mV/mm pulsed electric signal was applied to the scaffolds using an external pacing controller (C-Pace EP, IonOptix), at a frequency of 12 Hz, a pulse duration of 50 ms on a continuous application for 6 days. Metabolic activity was measured at days 1, 3 and 7 and following completion of the stimulation period, the scaffolds were washed with PBS and either fixed using 4% PFA for 1 hour for immunohistochemical analysis or frozen and stored for DNA quantification.

### 4.11 Assay analysis of cell growth

Metabolic Viability Assay Analysis: At selected time points an Alamar Blue assay (Invitrogen, UK) was performed according to the manufacturer’s instructions. For each time point, films, coverslips or scaffold samples were washed with DPBS and immersed in a 10% Alamar Blue, 90% medium solution (500 μL per well, 1 h, 37 °C). The assay solution was pipetted in triplicate into an opaque 96-well plate and the absorbance of the solution was read at 563 nm to give a relative comparison of cellular metabolic activity of the cells. The coverslips/scaffolds were then washed in DPBS and re-immersed in the appropriate culture medium.

DNA Quantification: DNA quantification was performed using a Quant-IT Picogreen dsDNA assay kit (Invitrogen, UK) according to the manufacturer’s instructions. Briefly, cells were lysed in 500 μL lysis buffer (0.2 m carbonate buffer containing 1% Triton-X100) and then underwent three freeze-thaw cycles at-80 °C to fully lyse all cells. Next, each sample was diluted 1:2 in 1× Tris-EDTA buffer before adding 100 μL of sample solution into an opaque 96-well plate in duplicate. Quant-iT dsDNA reagent was prepared by adding 100 μL to 19.9 mL of 1× Tris-EDTA buffer. Before adding 100 μL to each well-containing sample. The fluorescence of each sample was measured at an excitation wavelength of 485 nm and an emission wavelength of 538 nm. DNA concentration was calculated using a standard curve. Alamar blue and Picogreen results for each sample were used to calculate cell activity relative to cell DNA content.

### 4.12 Isolation and Characterization of Primary Olfactory Bulb-derived Stem Cells

Olfactory bulbs were dissected out of E18 – P2 wild-type mouse pups. The tissue washed in sterile PBS (4°C), before being digested in trypsin-EDTA (100mM) for 20 mins to release cells. Excess trypsin was pipetted from the cell suspension which was resuspended in ONSC seeding medium consisting of DMEM:F12 (1:1), 1% P/S, 1% Glutamax and 10% FBS and seeded in 6 well plates for 4 hrs (N=4-6 bulbs per well). All animals were kindly donated by fellow researchers at the Tissue Engineering Research Group in keeping with the 3Rs and the postmortem harvesting was carried out under HPRA individual license (AE19127/I259) and with ethical approval from the RCSI Research Ethics Committee (REC202005013).Once the primary cells attached, the medium was changed to ONSC growth medium consisting of DMEM:F12 (1:1), 1% P/S, 1% Glutamax, 2% B27, 20 ng/ml EGF, 20 ng/ml bFGF and 4 ug/ml porcine heparin sodium salt. Within 72 hrs, floating neurospheres were released from the adherent plastic surface which were transferred (2000 cells/well) to 96-well non-adherent Nunclon Sphera 96-well plates (Sigma) and allowed to form neurospheres. After 10 days the neurospheres were fixed and characterized using immunohistochemical analysis. Immunohistochemistry was used analyse the ONSC morphology and protein expression. Mycoplasma testing indicated no contamination.

### 4.13 Electrical Stimulation of ONSC-seeded MXene-ECM macroporous scaffolds

To assess the capacity of MXene-ECM scaffolds of varying conductive properties and microfibre densities, ONSCs were cultured for 1 week in non-adherent 96 well plates to form spheres before being seeded onto sterilized scaffolds in growth medium. After 24 hrs, the ONSC-laden scaffolds (n≥3) were transferred to 8 well plates and stimulated using the IonOptix Bioreactor, as previously described, for a further 6 days. The stimulation profile was 12 Hz, 200 mV/mm, 50 ms pulses. The scaffolds were then fixed in 4% PFA for 1 hr and stored at 4°C for immunohistochemical analysis.

### 4.14 Immunohistochemistry and Confocal Microscopy

Following fixation in 4% PFA (30 min, RT) and a DPBS wash step, astrocyte, microglia and neuron-seeded films, coverslips and scaffolds were permeabilized with 0.1% Triton X100– DPBS (30 min, RT), washed in DPBS before being incubated with blocking solution overnight at 4°C. Primary antibodies were then added as listed below and incubated overnight at 4°C. Samples were then washed in dPBS (x 3 times, 5 mins) before the addition of secondary antibody solution (listed below) and another overnight incubation step (4°C). Samples were washed and then incubated with Atto-Phalloidin (1:500, 2 h, RT) to highlight the actin skeleton before a further wash step and the incubation of samples with 4′,6-diamidino-2-phenylindole (DAPI) (1:1000, up to 1 h, RT) to label the cell nuclei prior to 3 final wash steps. Coverslips and films were then mounted on glass slides for imaging using Fluoromount mounting medium (Invitrogen, UK). Fluorescently labelled samples were imaged using a Zeiss 710 LSM NLO confocal microscope (Zeiss) at consistent exposure, gain, and magnification across all samples. The longest axon was measured manually using FIJI^[86]^ tools in each field of view on the neuron and neurosphere-seeded scaffolds and calculated as an average per sample. The average neurite length was measured using a FIJI plugin, Neurite Tracer^[87]^, to measure the total length of neurites in each image (4 images per scaffold, n ≥ 3 scaffolds per group). Protein expression was measured through analysis of the average pixel intensity of the target protein (e.g. βIII tubulin, GFAP) within the area of the cell outlined by phalloidin.

**Table 1.** Immunohistochemistry primary and secondary antibodies.

| Cell Type | Primary Antibody | Concentration | Manufacturer | Code |
| --- | --- | --- | --- | --- |
| Astrocytes | Mouse anti-Glial Fibrillary Acidic Protein | 1:500 | Sigma Aldrich, Ireland | 556328 |
| IMG Microglia | Rabbit Anti-IBA-1 | 1:500 | Abcam, UK | Ab5076 |
| SH-SY5Y Neurons | Rabbit Anti-βIII tubulin | 1:750 | Sigma Aldrich, Ireland | T2200 |
| SH-SY5Y Neurons | Mouse Anti-MAP2 | 1:500 | R&D Systems, MN, USA | MAB8304 |
| ONSCs, SH-SY5Y Neurons | Rabbit Anti-βIII tubulin | 1:750 | Sigma Aldrich, Ireland | T2200 |
| ONSCs | Mouse Anti-Glial Fibrillary Acidic Protein | 1:750 | R&D Systems, MN, USA | 556328 |
| ONSCs | Mouse Anti-Nestin | 1:500 | R&D Systems, MN, USA | MAB2736 |
| ONSCs | Mouse Anti-SOX2 | 1:500 | R&D Systems, MN, USA | AF3076 |
| Secondary Antibody |  |  |  |  |
| Goat anti-rabbit 488 |  | 1:1000 | Invitrogen, UK | A11036 |
| Donkey anti-mouse 555 |  | 1:1000 | Invitrogen, UK | A31570 |
| Goat Anti-mouse 647 |  | 1:750 | Invitrogen, UK | A21235 |

---

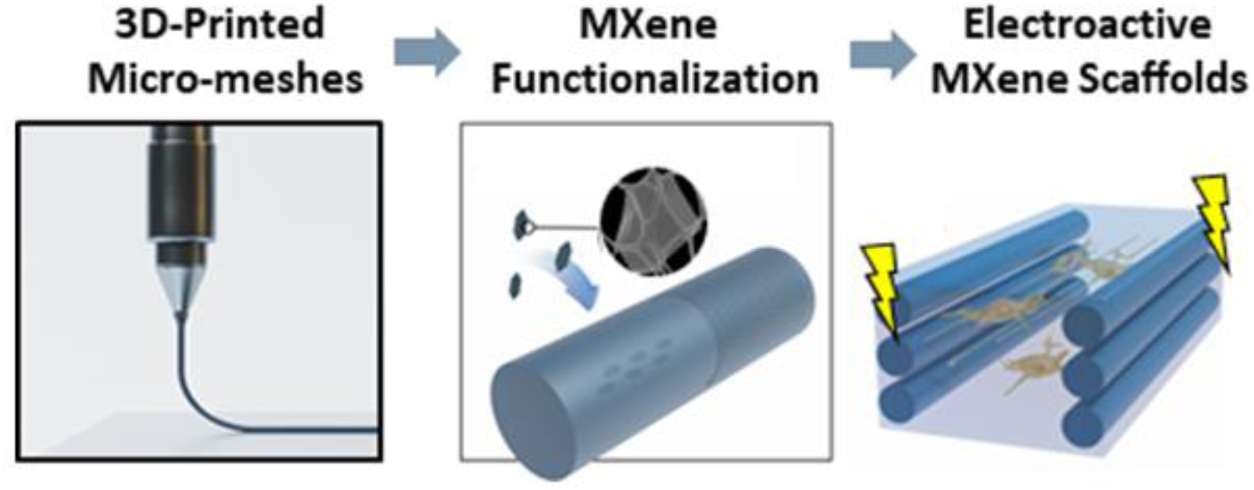

This study combined microscale 3D-printing with 2D Ti_3_C_2_T*_x_* MXene nanosheets to produce tunable electroconductive microfibrous micro-meshes. By embedding these MXene-functionalized micro-meshes within a growth-supportive macroporous extracellular matrix (ECM)-based biomaterial, a multifunctional biomimetic MXene-ECM scaffold was developed for neurotrauma repair. These MXene-ECM scaffolds enhanced the capacity of electrical stimulation to promote repair-critical processes in neuronal cells, in a manner dependent on electroconductive micro-mesh design.

