## Supplemental Figures for "3D-Printing of Electroconductive MXene-based Micro-meshes in a Biomimetic Hyaluronic Acid-based Scaffold Directs and Enhances Electrical Stimulation for Neural Repair Applications"

**Supplementary Figures:**

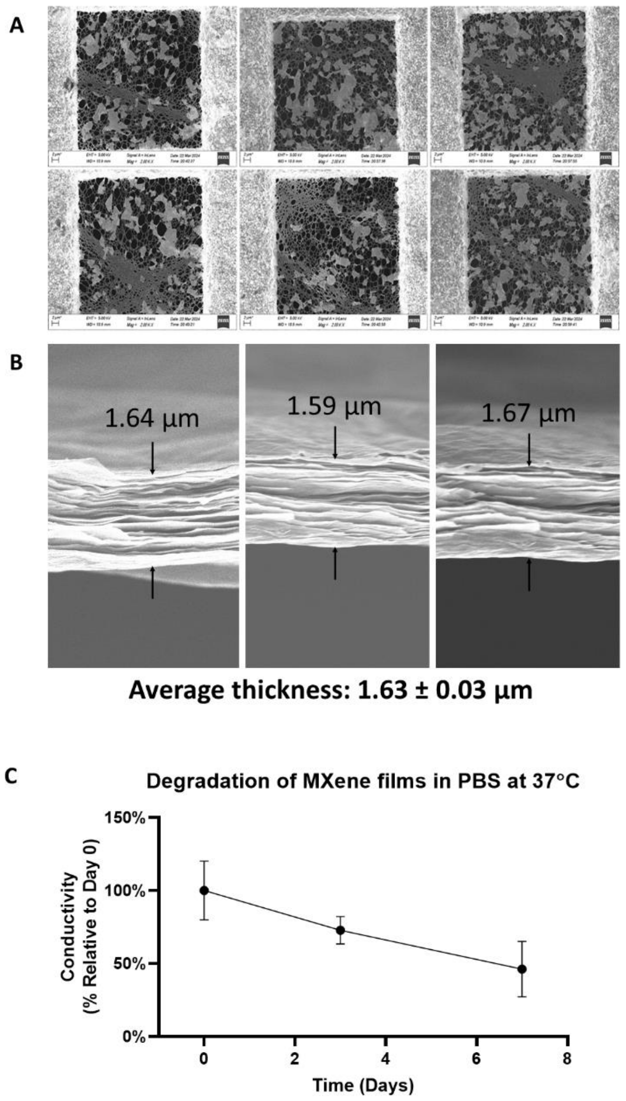


**Supplementary Figure 1.** MXene Characterization Data. (A) Size distribution of MXene nanosheets. (B) Cross-section of MXene films. (C) Degradation of electrical conductivity in ionic medium. MXenes exhibit robust degradation resistance over time in PBS salt solution mimicking the in vivo environment. MXene conductivity decreased to approximately 46.2 ± 19.1% after 7 days of degradation.

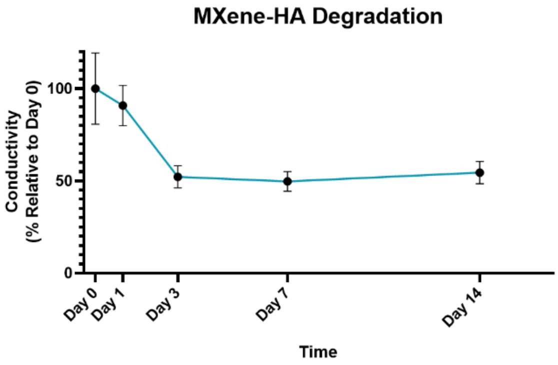


**Supplementary Figure 2. Degradation Characteristics.** Medium density MXene-ECM scaffolds incubated in PBS over 14 days. The MXene-ECM scaffolds exhibit an approximate 50% drop in conductivity over 3 days before stabilizing over the two week period. This degradation behaviour is most likely due to separation of the MXene flakes due to swelling and infiltration of water molecules.
